# A cell-free synthetic biochemistry platform for raspberry ketone production

**DOI:** 10.1101/202341

**Authors:** Simon J Moore, Tomasso Tosi, David Bell, Yonek B Hleba, Karen M Polizzi, Paul S Freemont

## Abstract

Cell-free synthetic biochemistry provides a green solution to replace traditional petroleum or agricultural based methods for production of fine chemicals. 4-(4-hydroxyphenyl)-butan-2-one, also known as raspberry ketone, is the major fragrance component of raspberry fruit and is utilised as a natural additive in the food and sports industry. Current industrial processing standards involve chemical extraction with a yield of 1-4 mg per kilo of fruit. As such its market price can fluctuate up to $20,000 per kg. Metabolic engineering approaches to synthesise this molecule by microbial fermentation have only resulted in low yields of up to 5 mg L^−1^. In contrast, cell-free synthetic biochemistry offers an intriguing compromise to the engineering constraints provided by the living cell. Using purified enzymes or a two-step semisynthetic route, an optimised pathway was formed for raspberry ketone synthesis leading up to 100% yield conversion. The semi-synthetic route is potentially scalable and cost-efficient for industrial synthesis of raspberry ketone.

## Introduction

For fine chemical biomanufacturing, synthetic biology aims to provide green solutions to replacing traditional petroleum or arable farming based production methods ^1–6^. The synthetic biochemistry approach, whereby metabolic pathways can be entirely reconstituted within a test-tube with purified enzymes ^7–11^ or cell-free extracts ^12–14^, offers a realistic concept to traditional cell based engineering, with the potential for high performance synthesis of fine chemicals and recombinant proteins to the industrial scale ^11,15^.

4-(4-hydroxyphenyl)butan-2-one, commonly referred to as raspberry ketone, is the major fragrance compound from raspberry berries (*Rubeus rubrum*). Raspberry ketone is harvested by chemical extraction at a ratio of 1-4 mg per kilo of fruit ^16^. This presents a highly-inefficient purification process, which is reflected by a high market price for naturally extracted raspberry ketone that fluctuates at approximately $10,000-20,000 kg^−1^, with global production estimated at between 100-200 tons per annum ^16,17^. As an alternative green solution for its production, raspberry ketone can be synthesised in microbial cells. However, current efforts in either *Escherichia coli* ^17^ or *Saccharomyces cerevisiae* ^17,18^ have only produced limited quantities of raspberry ketone (from 0.2 to 5 mg L^−1^), even despite the use of high cell-density fermentation. The raspberry ketone biosynthetic pathway in plants begins from tyrosine or phenylalanine and belongs to the flavonoid natural product family, which includes the medicinal compounds curcumin ^19,20^, naringenin ^21^, resveratrol ^22^ and gingerole ^23,24^. The synthesis of flavonoid natural products requires a dedicated type III polyketide synthase that uses the cofactor malonyl-CoA for chain extension. For raspberry ketone, the enzyme responsible for this step is the benzalacetone synthase (BAS), which converts *p*-coumoryl-CoA and malonyl-CoA into 4-hydroxybenzalacetone (HBA) using a unique decarboxylation event ^25^, thus affording the precursor of raspberry ketone. Possible limitations for engineering raspberry ketone in cells include a limited malonyl-CoA pool (~35 μM in glucose catabolism) in *E. coli* ^26,27^ and a comparatively slow turnover rate (0.1 s^−1^) for the BAS enzyme ^28^. Additionally, previous metabolic engineering studies have led to a mixed yield of raspberry ketone and its precursor HBA, which requires a further double bond reduction. Therefore, an efficient alkene reductase is also required to complete the pathway.

Routinely in synthetic biology, microbial cells are engineered through plasmid or chromosomal based heterologous gene expression to synthesise a desired synthetic fine chemical. However, this often presents a myriad of challenges to the living cell. This includes, but is not limited to, cellular burden ^29^, metabolic flux ^30^, accumulation of toxic intermediates ^4,31^, poor substrate availability ^26^ and non-productive chemical or enzyme based side-reactions ^32^. In addition, whilst *in vitro* deduced enzyme *k*_cat_ values are often reflective of *in vivo* catalytic rates (*k*_app_), it should also be considered that the cellular environment can modulate enzyme controlled pathways through changes in thermodynamics or substrate availability ^33^. The low yields of raspberry ketone obtained in microbial cells^17,18^ suggests that an enzyme bottleneck or product toxicity limits it’s production *in vivo*.

For fine chemical pathways, as well as drugs and recombinant proteins, an *in vitro* approach offers a unique opportunity to the biochemist to control and modify synthetic pathways outside the regulatory control of the cell. Using raspberry ketone as a model pathway, we detail a purified enzyme approach to achieve a high-yield synthesis in a one-pot reaction. Together with other recent synthetic biochemistry studies ^11,13^, we feel this approach is potentially expandable to other high-value fine chemicals that require cost efficient cofactor regeneration (i.e. malonyl-CoA, ATP) for *in vitro* synthesis. In summary, we present a modularised enzyme prototyping approach to completely convert the substrate tyrosine via a five-enzyme cascade into raspberry ketone, as well as demonstrating how enzyme levels competing for the key cofactor, coenzyme A (CoA), require balancing for optimal performance. To complement these findings, we also present a crystal structure of the NADPH-dependent raspberry ketone reductase, which we use to relax its cofactor specificity towards NADH utilisation for an improved low-cost *in vitro* engineering.

## Results

### Enzyme synthesis of raspberry ketone pathway *in vitro*

In order to begin studying the raspberry ketone pathway *in vitro* and identify key metabolic bottlenecks, we selected a synthetic five-step pathway for raspberry ketone using a combination of bacterial, fungi and plant enzymes (Figure 1A). The starting point for this pathway begins from tyrosine, the natural substrate for raspberry ketone synthesis in plants, using tyrosine ammonia lyase (TAL) to deaminate tyrosine, forming *p*-coumarate ^34^. This in turn is activated for malonyl-CoA extension by the addition of CoA by an ATP-dependent CoA ligase (PCL) to form *p*-coumoroyl-CoA ^35^. Following this, the benzalacetone synthase (BAS) uses malonyl-CoA for chain extension via a double decarboxylation event to release the HBA product ^25,28^. Finally, a NADPH-dependent raspberry ketone synthase (RKS), a double bond reductase, converts HBA into raspberry ketone ^36^. Based on this synthetic pathway, a set of highly active and kinetically well-characterised enzymes for TAL, PCL and a malonyl-CoA synthetase (MatB) were selected from the BRENDA database (see materials and methods), together with BAS and RKS (previously abbreviated as RZS for raspberry ketone/<u>z</u>ingerone <u>s</u>ynthase), where only a single enzyme has been reported and characterised ^25,36^. Initially, each enzyme was recombinantly produced in *E. coli* BL21 Gold (DE3) pLysS and purified at high-yields (~10-100 mg L^−1^) from the soluble fraction with at least 95% purity as estimated by SDS-PAGE (Figure 1B). The activity of TAL and PCL enzymes was also assayed individually (Supplementary Figure S1) and results were in close agreement to previous reported literature values ^37,38^.

**Figure 1.**
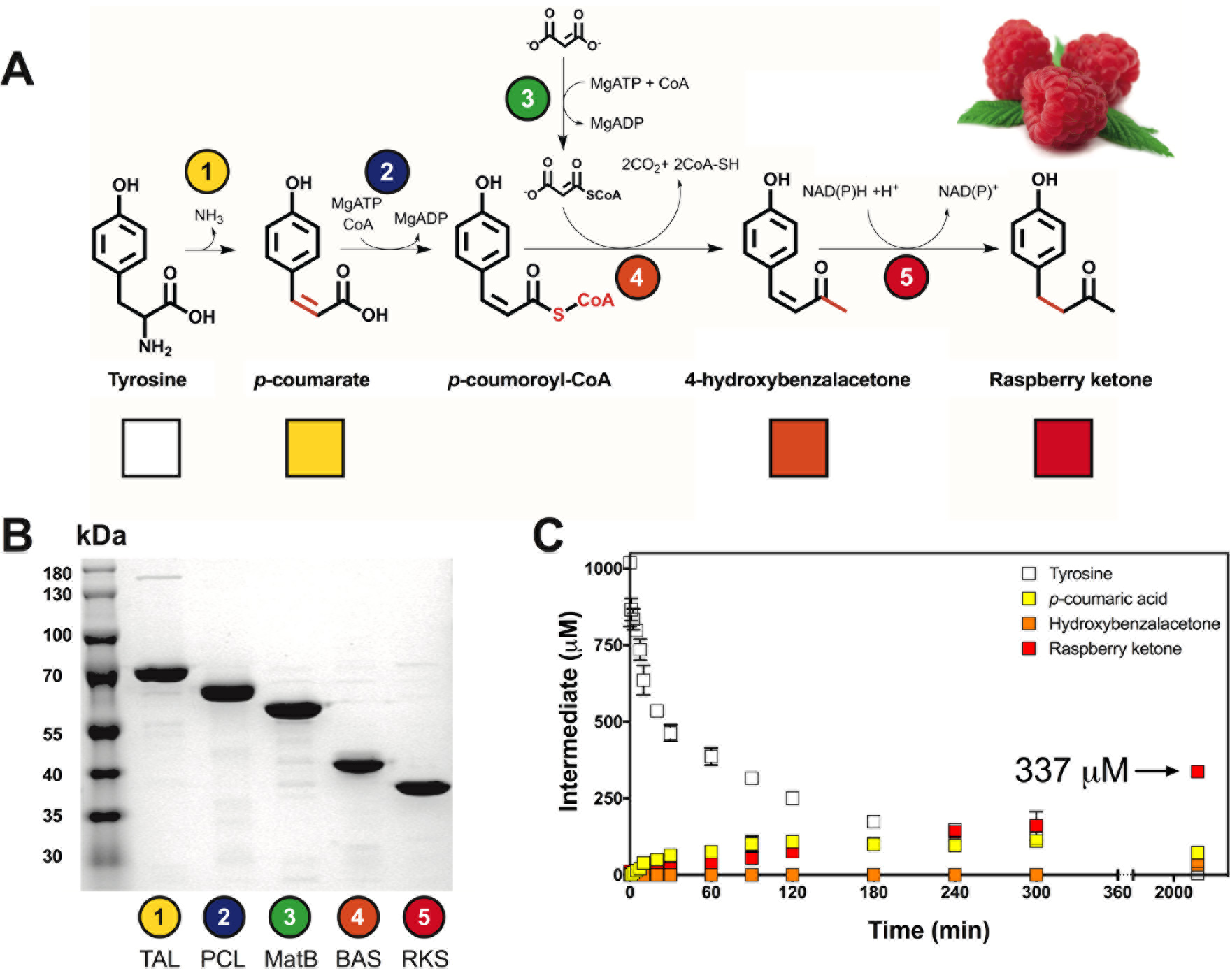
A synthetic biochemistry module for raspberry ketone synthesis *in vitro*. (A) A synthetic pathway for raspberry ketone synthesis from Tyrosine using malonate for malonyl-CoA synthesis. Intermediates quantified by LC-MS include Tyrosine (white box), *p*-coumarate (yellow box), hydroxyHBA (orange box) and raspberry ketone (red box). (B) 2 μg of purified enzyme was loaded and analysed by 12% SDS-PAGE and Coomassie Blue staining. Sizes of His6-tagged recombinant enzyme - *R. glutinis* TAL (77.0 kDa), *A. thaliana* PCL (63.2 kDa), *R. palustris* MatB (56.6 kDa), *R. palmatum* BAS (44.4 kDa), *R. rubrum* RKS (40.7 kDa). (C) One-pot synthesis of raspberry ketone with 2.5 μM of enzymes. For full details, please refer to Supplementary Table S2.

To test the synthetic *in vitro* pathway, 5 μM of purified TAL, PCL, MatB, BAS and RKS enzymes were incubated with 1 mM tyrosine and substrates/cofactors (MgATP, CoA, malonate and NADPH). Please refer to the materials and methods for further details. Samples were removed at time points and quantified by reverse phase (C18) liquid-chromatography mass spectrometry (LC-MS) (Figure 1C). To begin, tyrosine is steadily depleted, whilst *p*-coumarate accumulates up to a maximum of 110 μM. Under these conditions, the HBA intermediate is not detectable, with the RKS fully converting its substrate into raspberry ketone. Raspberry ketone synthesis occurred at an initial linear rate of 0.63 μM min^−1^ for up to 6 hours, with a 33.7 % yield (337 μM) achieved at end-point, which is equivalent to 55 mg L^−1^ under batch synthesis. This suggested that the synthetic pathway was highly active in relative comparison to previous low yields achieved from cell-based metabolic engineering efforts (from 0.28 mg L^−1^ and 5 mg L^−1^ in yeast and *E. coli*, respectively) ^17,18^.

### A fluorescence biosensor for optimising flavonoid synthesis *in vitro*

Following the initial demonstration of raspberry ketone synthesis *in vitro*, we next sought to improve the overall pathway performance by identifying key bottlenecks and optimal pathway conditions. Due to the presence of an extra double bond, the product of the BAS enzyme, HBA, shares broad UV-Visible absorbance with the precursor intermediates *p*-coumarate and *p*-coumoroyl-CoA. Therefore, it was not possible to rapidly optimise the pathway with respect to BAS activity using a UV-Visible absorbance assay. Instead, the pathway was modularised by designing a fluorescence-based product detection assay to separately optimise *p*-coumoryl-CoA and malonyl-CoA synthesis, the shared substrates for the BAS enzyme (Figure 2A) with the rationale that a delicate balance of these is likely necessary to maximise total production concentration because they both rely on CoA for their synthesis.

**Figure 2.**
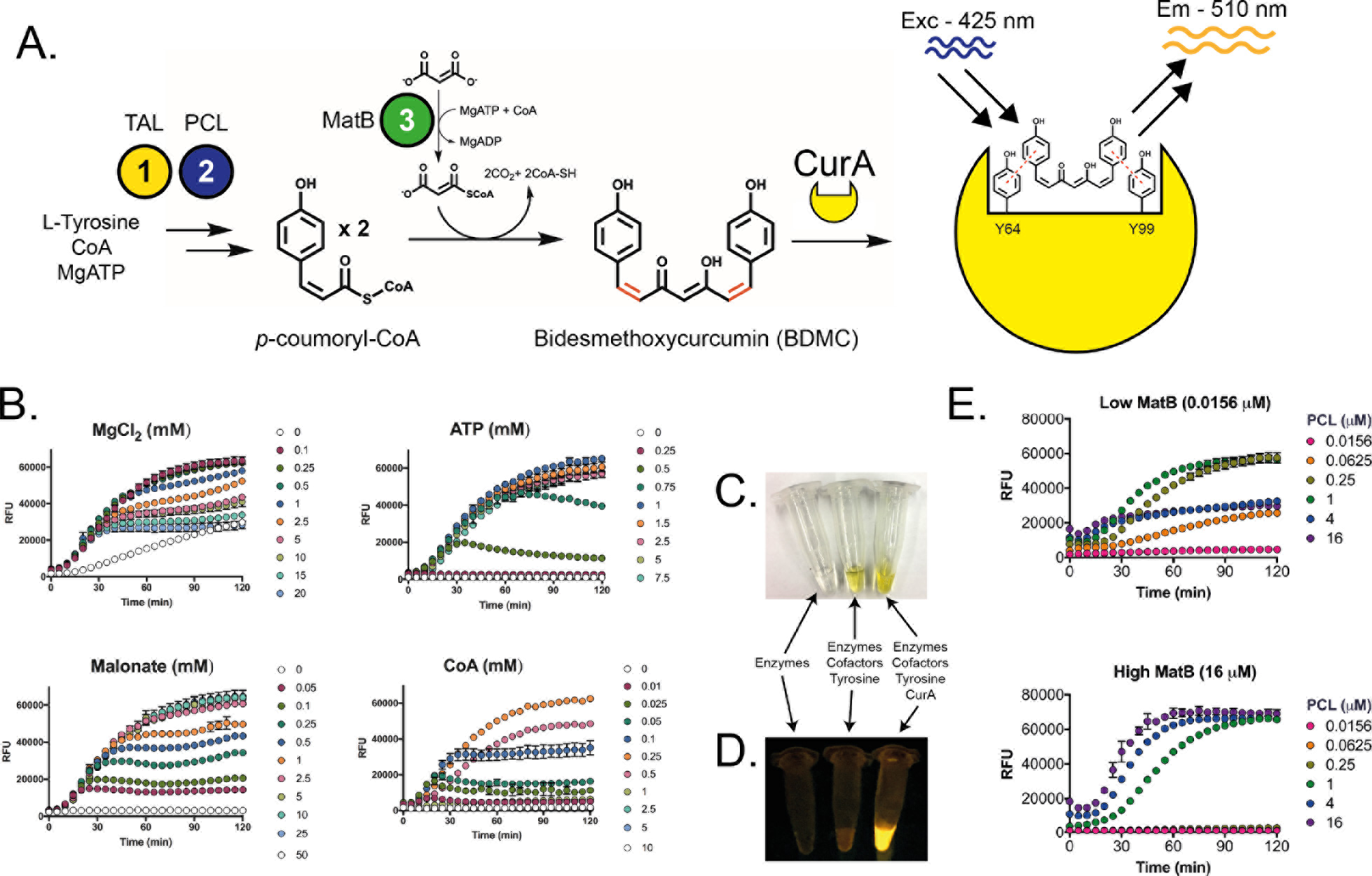
Development of a fluorescence sensor for type III polyketide synthesis. (A) A synthetic pathway for detection of BDMC synthesis from malonyl-CoA and *p*-coumoroyl-CoA. CurA a BDMC/curcumin NADPH-dependent reductase from *E. coli* is used in the absence of NADPH to bind to BDMC/curcumin generating an unique fluorescent output for relative quantitation of pathway activity. (B) Optimisation of cofactors and substrates for type III polyketide synthesis. (C) Visual and (D) fluorescence of *in vitro* BDMC reactions, with negative controls. (E) Enzyme competition between MatB and PCL, which share CoA and ATP for activity.

Previously, curcumin, which is a structural dimer of raspberry ketone, was identified to exhibit fluorescence when bound non-specifically to Gram-negative bacterial extracellular curli (amyloid-type) protein fibres ^39^. The precursor of curcumin, bidesmethoxycurcumin (BDMC), is synthesised from *p*-coumoroyl-CoA and malonyl-CoA by the curcumin synthase (CUS) ^40^, a type III polyketide synthase that is related to the BAS enzyme with 69% amino acid identity. Additionally, a NADPH-dependent curcumin double bond reductase (CurA), which is related to the RKS reductase enzyme with 38% amino acid identity, was previously characterised from *E. coli* ^41^. A Phyre2 ^42^ structural model of CurA suggested a large active site dominated by Tyr and Phe residues. Based on the potential fluorescence properties of non-specifically bound curcumin, we speculated that substrate binding of the apo-form of CurA to either curcumin or BDMC would also generate fluorescence through *π*-*π* stacking within the aromatic active site. To test this, we overexpressed and purified CurA with an N-terminal His_6_-tag from *E. coli* BL21 Gold (DE3) pLysS. Mixing an equal amount (10 μM) of apo-CurA with curcumin resulted in strong fluorescence with a maximum excitation and emission peaks at 425 nm and 520 nm, respectively, indicating ligand binding (Supplementary Figure S2). If an excess (100 μM) of NADPH was also added, the yellow coloured curcumin was rapidly reduced into the colourless tetrahydrocurcumin. This resulted in a quenching of the fluorescence, due to double bond reduction and a loss of conjugation. This suggested that CurA could bind curcumin in the absence of NADPH and therefore does not require the formation of a ternary complex. Next, with 25 μM of CurA incubated with an increasing concentration of curcumin (0-100 μM), the fluorescence signal fitted an exponential saturation curve (Supplementary Figure S3). Further increases in signal could be achieved with higher levels of CurA protein, but since the yield of recombinant CurA in *E. coli* was limiting (~10 mg L^−1^), a maximum level of 25 μM was used for subsequent assays to preserve protein stocks.

To confirm the use of this protein-ligand sensor as a real-time fluorescence assay of BDMC/curcumin synthesis, 25 μM of CurA was mixed in a one-pot reaction containing 1 μM of TAL, PCL, MatB and the type III polyketide synthase, CUS, along with the necessary cofactors and 1 mM Tyrosine. In addition, each enzyme was individually omitted as a control to confirm the specificity of the assay. Following a time-course reaction, an increasing fluorescence signal was observed (Supplementary Figure S4), relative to a stable background signal if any of the enzymes or substrates were omitted (Supplementary Figure S4). The initial rate of the fluorescence signal was further improved by increasing the concentration of the substrate tyrosine or the CUS enzyme (Supplementary Figure S5). Furthermore, if a 1 μM aliquot of NADPH was injected midway through the reaction, a reduced fluorescence was observed, confirming that reduction of the double bond quenches the fluorescence. This confirmed the sensitivity and specificity of the assay for BDMC synthesis.

### Optimising the synthesis the substrates *p*-coumoryl-CoA and malonyl-CoA

To potentially improve the activity of the BAS enzyme, we next sought to use our novel fluorescence small molecule sensor to rapidly optimise the synthesis of the substrates *p*-coumoryl-CoA and malonyl-CoA *in vitro*. Firstly, one-pot synthesis of BDMC accumulated a yellow visual appearance (Figure 2C), which was fluorescent in the presence of the CurA sensor (Figure 2D). Initially, the conditions for *in vitro* synthesis of BDMC were optimised. Firstly, tyrosine and CUS levels were set at 5 mM and 25 μM, respectively, so that these parameters were not rate limiting (Supplementary Figure S5). An important consideration for *in vitro* enzyme systems is the concentration of substrates and cofactors (Figure 2B). BDMC synthesis was next optimised with respect to ATP, Mg^2+^, CoA and malonate levels. As expected, the ATP level was critical to BDMC synthesis with at least 1 mM required to reach fluorescence saturation (~65,000 RFU), whilst for optimal malonyl-CoA supply, a concentration of greater than 2.5 mM malonate was required for maximal activity (>60,000 RFU) and fluorescence saturation. The Mg^2+^ levels were equally optimal at 0.1-1 mM, but with higher levels, a biphasic fluorescence curve was observed (Figure 2B). A similar observation also occurred with the levels of CoA, where concentrations below the optimum level (0.25 mM) were also found to demonstrate a biphasic response. One potential interpretation of this response is a temporary depletion of free CoA availability, thus leading to an imbalance between *p*-coumoryl-CoA and malonyl-CoA levels for the CUS enzyme. In contrast, if higher levels of CoA (>1 mM) were added, complete inhibition of CurA-BDMC was observed (Figure 2B).

We next used this optimised set of conditions to determine if the enzyme levels of TAL, PCL or MatB control BDMC synthesis. To do this, each enzyme was varied in a 4-fold dilution series from 0.0156 to 16 μM. Firstly, some background fluorescence attributed to high levels of *p*-coumoryl-CoA occurred if the level of the PCL enzyme was increased above 1 μM (Figure 2E). However, if both the levels of the TAL and PCL enzymes were optimised to a peak concentration of 16 μM and 1 μM, respectively, maximal BDMC synthesis (87,400 RFU) was reached. Interestingly, by varying PCL and MatB together, noticeably a careful balance of enzymes was required to prevent pathway inhibition, possibly through depletion of free CoA (Figure 2D). This evidence overall suggests that the pathway is tightly regulated by a sensitive interplay between the free CoA pool, CoA derivatives, CoA-dependent enzymes and the rate-limiting activity of the type III polyketide synthase (CUS). Therefore, whilst this experiment only provides a relative measure of pathway activity, it demonstrates that a finely tuned interplay between enzyme levels and the CoA cofactor is required for optimal pathway performance. This observation would not be possible to detect through cell-based engineering. Based on these measurements, we next used the optimal parameters to determine whether raspberry ketone synthesis could be improved.

### Raspberry ketone biosynthesis is inhibited by a high tyrosine concentration

We next sought to increase the yield of raspberry ketone synthesis *in vitro* by providing optimised levels of *p*-coumoroyl-CoA and malonyl-CoA synthesis using the parameters determined in CurA-BDMC fluorescence assay. Firstly, batch reactions were prepared with varying concentrations of tyrosine from 100-1000 μM. Between 100-300 μM tyrosine, raspberry ketone was produced at a 100% yield (Figure 3A). Beyond this, the level of *p*-coumarate rises, without a further increase in raspberry ketone yield. Initially, we suspected that the supply of malonyl-CoA or *p*-coumoryl-CoA was limited by the exhaustion of the ATP supply. However, although providing ATP regeneration through phosphoenolpyruvate (PEP) and pyruvate kinase increased the flux of tyrosine and *p*-coumarate, the net yield of raspberry ketone was lowered (Figure 3B). Unexpectedly, these conditions led to an accumulation (~30-50%) of the BAS catalysed side-product bisnoryanogonin (BNY), in comparison to standard conditions where only minor levels (~5%) were detected by LC-MS. BNY is a side-product of the BAS enzyme that is produced by an additional malonyl-CoA extension of the diketide intermediate ^43^. Previously, BNY was produced only at minor levels at mildly alkaline pH (7.5-8.5), but elevated at pH 6.0-7.5 in Tris-HCl or phosphate buffers ^43^. To attempt to correct this issue, we repeated the synthesis of raspberry ketone in a range of alkaline buffers (pH 7.5-9.0) and as a time-course at pH 9 (Supplementary Figure S7). Increasing the pH favoured the activities of the TAL and PCL enzymes, but this did not further increase the levels of raspberry ketone and only a minor decrease in the levels of the BNY side-product was observed. Under these forced synthetic conditions, it appears that the HBA synthesis activity of the BAS enzyme switches towards a preference of BNY synthesis. Since we could not find a route to eliminate BNY synthesis and this capped our overall yield of raspberry ketone, we next considered how the cost-efficiency of this *in vitro* pathway could be improved by relaxing the cofactor specificity of the NADPH-dependent RKS reductase step.

**Figure 3.**
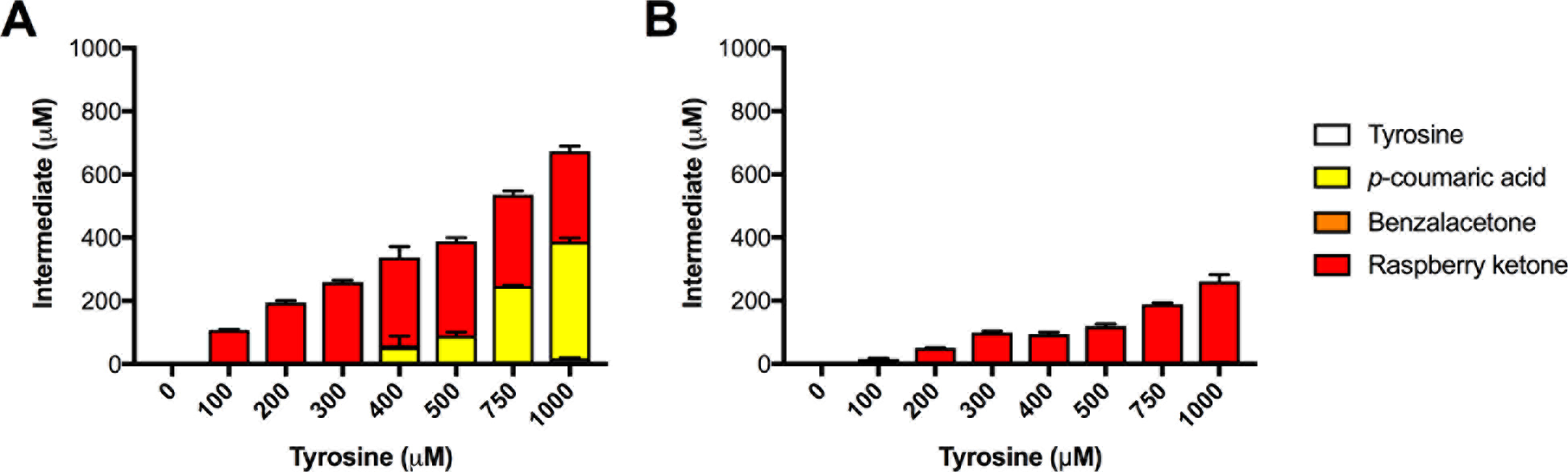
One-pot synthesis of raspberry ketone with a varying concentration of tyrosine. (A) Standard conditions and (B) optimised conditions as outlined in Supplementary Table S2.

### NADH is rate limiting for *in vitro* raspberry ketone synthesis

The final step of raspberry ketone synthesis is catalysed by the NADPH-dependent double bond reductase RKS. For *in vitro* based biocatalysis, NADH ^44^ or biomimetic analogues ^45^ are preferred to provide increased stability and reduced cost for reductive enzyme reactions. We therefore revisited the one-pot synthesis of raspberry ketone providing NADH, as opposed to NADPH, as a cofactor (Supplementary Figure S6). With this, whilst the rate of tyrosine deamination by TAL remains unchanged, an increased accumulation of *p*-coumarate (172 μM) and HBA (15 μM) is observed after 6 hours, with only trace levels of raspberry ketone detected. Unsurprisingly, only trace levels of raspberry ketone were detected during the time-course, whilst after 48 hrs, a 7.6% (76.1 μM) yield of raspberry ketone was achieved. The initial lag in raspberry ketone synthesis confirmed that the RKS enzyme had a clear preference for NADPH as a cofactor.

To further understand the RKS reductase, the enzyme was next characterised *in vitro*. Firstly, HBA the natural substrate for RKS is yellow in colouration at pH >6 and strongly absorbs between 250-450 nm, which overlaps with NAD(P)H absorbance at 340 nm for kinetic characterisation. Instead, to initially obtain kinetic parameters for the RKS enzyme, the substrate analogue phenylbuten-2-one was used since it lacks UV-Visible absorbance at 340 nm. To begin, the RKS enzyme shares sequence similarity (77% amino acid identity) and similar kinetic properties to the previously characterised promiscuous *Nicotiana tabacum* double bond reductase NtDBR ^46^. For example, both RKS and NtDBR share increased activity towards acidic pH conditions (Supplementary Figure S8). For all further RKS enzyme assays, these were measured at 30°C and pH 6.4, with 1 mM of phenylbuten-2-one. Under these conditions, RKS shows an apparent *K*_cat_/*K*_m_ of 58 and 1.3 mM s^−1^ mM^−1^ with NADPH and NADH, respectively (Supplementary Table S3). This demonstrates a 45-fold preference for NADPH as a cofactor.

### Structure guided engineering of a NADH proficient RKS

A number of studies have highlighted that the cofactor specificity of NADPH-dependent reductase enzymes can be relaxed towards NADH utilisation ^47–50^ by modulating the interaction of the enzyme from the ribose 5’-phosphate (NADPH) in preference to the ribose 5’-hydroxyl group (NADH) ^51,52^. To aid in the rational design of increasing NADH activity of the RKS enzyme, we crystallised and solved the tertiary structure complex of RKS with raspberry ketone pathway substrate HBA (Figure 4A) and the cofactor NADPH (Figure 4B), deposited as pdb: 6EOW. By analysing the 2*F*_*0*_ – *F*_*c*_ Fourier syntheses, two configuration states were observed at a 50:50 ratio due to crystal soaking. The mixed states could be as a result of NADPH and ternary complex formation to the active site, thus displacing any NADP^+^ present from crystallisation. In the ternary complex state, additional electron density for HBA binding is observed, with *π*-*π* stacking between the HBA and nicotinamide aromatic rings (Figure 4A) with a hydride transfer distance of 3.06 Å to the alkene double bond. Furthermore, in the ternary complex, an additional patch of electron density consistent of a flexible loop supporting Y72 is observed. This flexible loop forms a cap over the active site (closed loop), with the *para* hydroxyl group in HBA moving towards Y72. With respect to HBA binding, the substrate is bent within the active site and is encased by aromatic residues Y59, Y72, Y85, F107, Y263 and F290 (Supplementary Figure S9). We speculate that this this closed loop formation holds the HBA within the active site prior to transition state. Interestingly, in contrast to apo-RKS and NtDBR, the Y72 flexible loop lacks electron density and seemingly points away from the active site (open loop state). Finally, in regards to cofactor binding, in both states the NADP(H/^+^) cofactor is bound in a typical conformation as is observed in the previous related structures ^46,53^ with the binding specificity provided by a triad of hydrogen bond based contacts with the G191 backbone nitrogen and the side chains K195 and Y211 (Figure 4B). Whilst K212 is also present, it is rather removed from binding to the ribose 5’-phosphate. Importantly, K195 can interact with either the ribose 5’-phosphate or neighbouring 4’-hydroxyl group.

**Figure 4.**
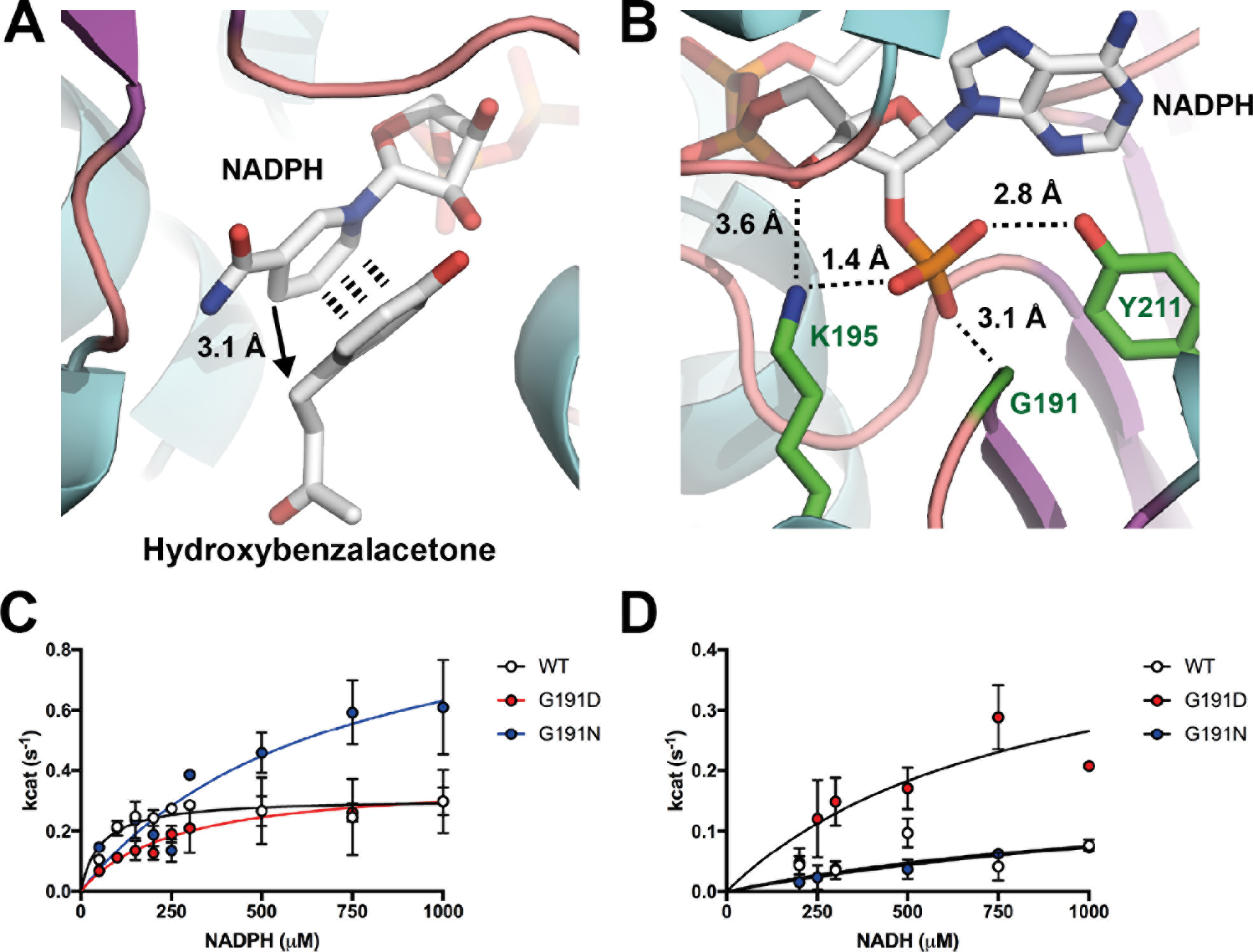
Structure based engineering of the RKS reductase towards NADH utilisation (pdb: 6EOW). (A) Binding site for HBA in proximity to the NADPH cofactor. (B) Cofactor specificity is provided by a triad of binding residues G191, K195 and Y211 with hydrogen bonding to the 5’-ribose phosphate. (C) Kinetic characterisation with NADPH and RKS variants and with (D) NADH as outlined in materials and methods.

In an effort to modify the cofactor specificity of the RKS enzyme, a number of amino acid substitutions at G191 and Y211 were made and steady-state kinetics was determined where appropriate (Figure 4C and Supplementary Table S3). We did not attempt to modify K195 due to its role in bonding to the 4’ hydroxyl group found in both NADPH and NADH. We instead chose to modify G191 to provide a short polar side chain (D and N) that can provide a hydrogen bond for the 5’-hydroxyl group unique to NADH. Both the D191 and N191 modifications decreased the *k*_cat_/*K*_m_ from 58 mM^−1^ s^−1^ (RKS^wt^) with NADPH to 14 and 15 (s^−1^ mM^−1^), respectively (Figure 4C]]). This decrease in catalytic efficiency was due to an increased *K*_m_ for NADPH with the variants. Interestingly, for N191, the *k*_cat_ increased by 3-fold up to 1.07 s^−1^ in comparison to the wild-type (0.31 s^−1^). One possible interpretation is that steric hindrance of N191 with the 5’ phosphate increases the catalytic rate by enhancing NADP^+^ release through decreasing its binding affinity. In contrast, with NADH as the cofactor, whilst N191 did not improve NADH utilisation, D191 was found to increase the *k*_cat_/*K*_m_ from 1.3 mM^−1^ s^−1^ (RKS^WT^) to 5.9 mM^−1^ s^−1^ (Figure 4D and Supplementary Table S3). In summary, these mutagenesis experiments at the G191 position demonstrated that the negatively charged (D) side chain improves NADH specificity in preference to a positive charged residue (N). Whilst further modifications to the Y211 position were also tested individually or in combination with D191 and N191 (data not shown), no further improvements in NADH specificity or *k*_cat_ were found, with most modifications leading to a loss of activity with either NADH or NADPH.

### NADH based *in vitro* synthesis of raspberry ketone from HBA

Whilst the kinetics of RKS (wild-type and D191) were not characterised with its natural substrate, HBA, its activity was demonstrated with NAD(P)H cofactor regeneration. Firstly, HBA was synthesised from the inexpensive substrates *p*-benzaldehyde ($203 per kg) and acetic acid ($67 per kg) using an aldol condensation reaction under basic conditions to provide a 76% yield ^54^ (Figure 5A). Then, using the thermostable phosphite dehydrogenase (PtxD) mutant Opt12 ^55^ for NAD(P)H regeneration from phosphite, we tested the reduction of the HBA substrate at 30°C by following a loss of absorbance at 400 nm (A400). For negative controls, in the absence of the reductase, phosphite or PtxD, the A400 for 1 mM HBA remained stable over the time-course measured (Figure 5B). With 1 mM HBA, 10μM of RKS, 20 mM phosphite and an excess of PtxD opt12, complete reduction was achieved with as little as 10 μM of NADPH (Figure 5B). Next, to demonstrate the proficiency of the RKS variants (10 μM) with both NADPH and NADH, a time-course reaction was monitored with 1 mM injections of HBA every hour with 0.25 mM NAD(P)H, 20 mM phosphite and an excess of PtxD opt12 (Figure 5B). For RKS^WT^, an initial rate of 59.1 and 12.8 μM/min/mg was observed with NADPH and NADH, respectively (Supplementary Table S2). In comparison, for RKS^D191^, the rates of reduction with NADPH and NADH were nearly equivalent at 46.5 and 43.5 μM/min/mg, respectively (Supplementary Table S2). This demonstrated high-yield and efficient synthesis of raspberry ketone with the RKS^D191^ variant providing an elevated and complete turnover (~100%) with the inexpensive NADH cofactor. RKS reductase activity was also stable for several days at room temperature (data not shown).

**Figure 5.**
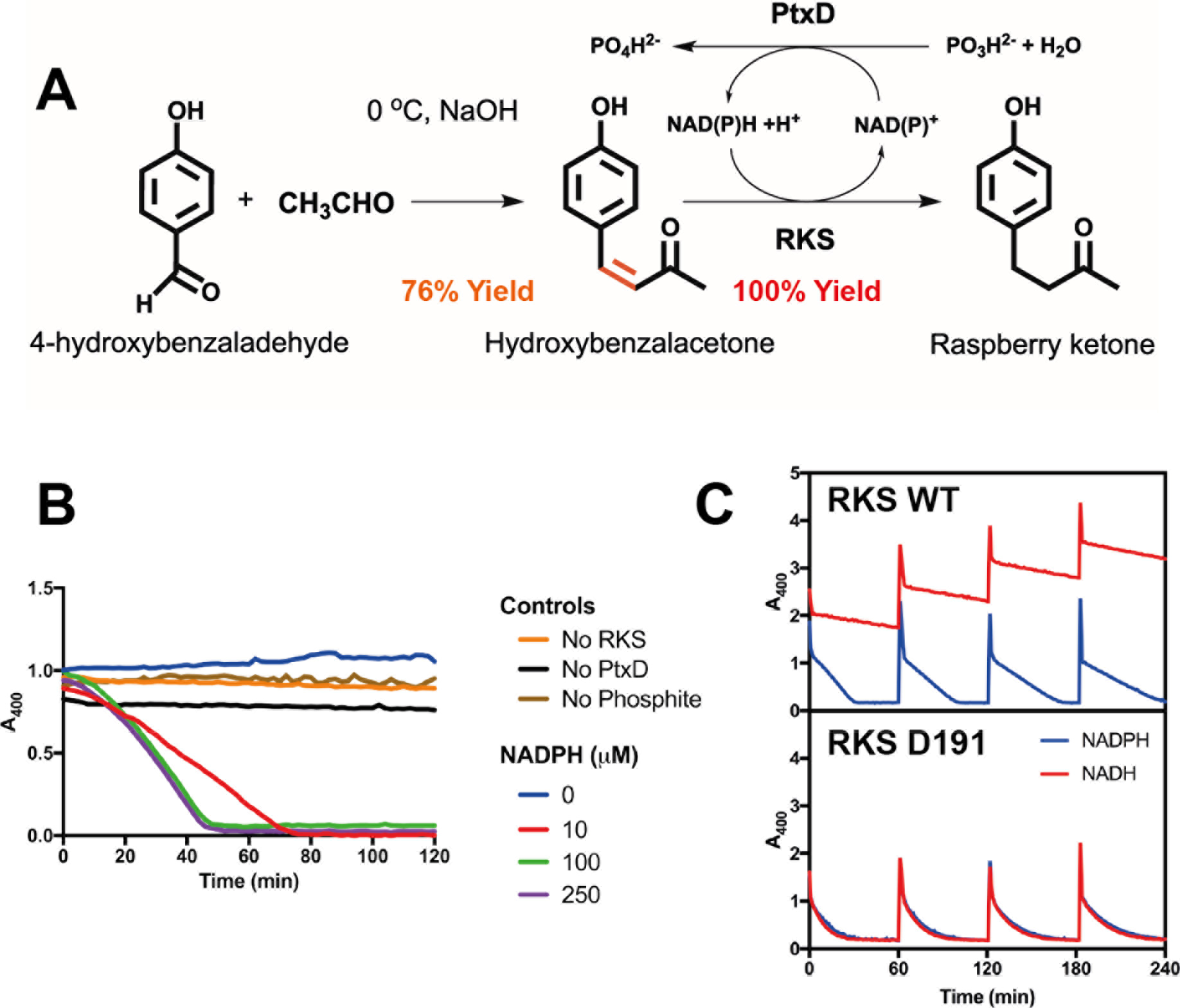
A two-step semi-synthetic route to high-yield raspberry ketone synthesis with NADH and cofactor regeneration. (A) A semi-synthetic pathway for raspberry ketone using aldol condensation and RKS reductase activity with cofactor regeneration. (B) Activity of the thermostable PtxD opt12 phosphite dehydrogenase with 1 mM HBA and 10 μM RKS. Negative controls (no enzyme or phosphite) are shown along with a variable concentration of NADPH. (C) Time-course reaction monitoring loss of absorbance at 400 nm showing reduction of HBA to raspberry ketone. Injections of 1 mM HBA were added in 60 min cycles. An excess of PtxD and 20 mM phosphite was incubated at 30°C with 10 μM RKS (top panel) or the D191 variant (bottom panel), with either 0.25 μM NADPH (blue line) or NADH (red line).

### NADH based *in vitro* synthesis of raspberry ketone from tyrosine

To understand whether the RKS^D191^ variant could be used for *in vitro* synthesis of raspberry ketone with NADH, three time-course reactions were prepared with 1 mM tyrosine and optimised enzyme levels and cofactors with either an absence of the reductase or a combination of RKS^WT^/NADPH or RKS^D191^/NADH. Firstly, ATP regeneration was omitted to limit the accumulation of the BNY intermediate. Then, without the RKS reductase, the reaction accumulated HBA at a rate of 0.81 min^−1^ (Supplementary Table S2), with a maximum yield of 421 μM observed after 48 hrs, along with 197 μM of leftover *p*-coumarate. By adding 20 μM RKS^WT^ with 0.5 mM NADPH along with cofactor recycling with the phosphite dehydrogenase ^55^, in contrast, a linear rate of raspberry ketone synthesis of 0.38 μM min^−1^ was observed, whilst after 48 hrs, the final concentrations of the intermediates were 291 μM *p*-coumarate and 375 μM raspberry ketone, with tyrosine or HBA undetected (Figure 6A). The HPLC chromatogram trace suggested that the remaining mixture was composed of the BNY side product. Finally, with 20 μM RKS^D191^ introduced with 0.5 mM of NADH and the recycling enzymes, in contrast to its earlier use with the wild-type RKS, the final level of raspberry ketone synthesis was improved to 297 μM at a rate of 0.51 μM min^−1^. However, a mixture of 297 μM *p*-coumarate and 198 μM HBA still remained present (Figure 6B).

**Figure 6.**
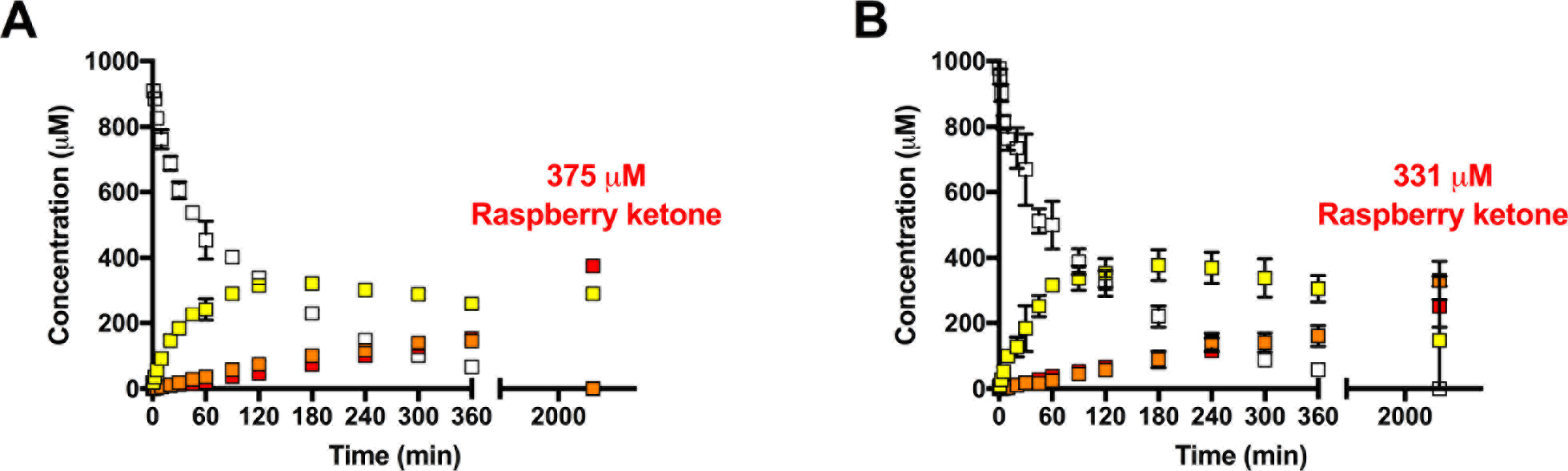
One-pot synthesis of raspberry ketone under optimised enzyme levels and cofactor regeneration. (A) Reaction with RKS^WT^ and NADPH. (B) Reaction with RKS^D191^ and NADH. Full synthesis conditions are provided in Supplementary Table S2.

## Discussion

Synthetic biochemistry offers an exciting opportunity to design and engineer enzyme pathways outside of the constraints of a living cell. Recent advances include the generation of a synthetic CO_2_ fixation cycle ^56^ and the potential industrial scale synthesis of bioplastic ^11^. Here, we show the application of the synthetic biochemistry approach to the synthesis of the natural product raspberry ketone. There is increasing interest in natural product biosynthesis as a source of new antimicrobials, human therapeutics, and fine chemicals. The synthetic biochemistry platform, which allows the rapid optimisation of pathway flux ^12,57^ using purified enzymes from different sources to identify high yielding combinations, is a powerful tool for natural product biosynthesis. Potentially, it could be used as a drug discovery tool, e.g. for probing the function of newly identified gene clusters in a sequential manner that allows the documentation of individual enzyme activities.

Natural products and other high-value fine chemicals are often only synthesized at very low concentrations in their natural source. Unless chemical logic (i.e reactive species) dictates, metabolic pathway enzymes lack selective pressure to evolve enhanced catalytic rates. For raspberry ketone, only tiny quantities are synthesised in the berries themselves and the process requires weeks of maturation. Therefore, yields as low as 1-4 mg per kilo of raspberry are obtained by chemical extraction of natural raspberry ketone fragrance ^17^. To meet the demand of raspberry ketone for the food and sports industry, a two-step organic synthesis via an aldol condensation followed by a rhodium-catalysed hydrogenation reaction is used ^58^. However, there is interest in developing a greener approach using synthetic biology to obtain this natural product. Previous work has investigated synthetically engineered *E. coli* ^17^ and Baker’s Yeast ^17,18^ systems, but yields from fermentation have been limited.

One of the advantages of the synthetic biochemistry approach is the ability to produce compounds that are toxic to cells or require high levels of enzyme expression to overcome pathway bottlenecks, which can lead to metabolic burden *in vivo*. Processes can also be designed to convert central metabolites to products without syphoning off essential resources required for cellular growth. Metabolic engineering *in vivo* requires a delicate balance between flux towards the product and towards biomass accumulation. When flux towards the product is sufficiently high to reduce cellular growth rates, there is inherent evolutionary pressure to lower productivity because mutations that reduce enzyme expression or primary metabolite usage will lead to a selective advantage. Our synthetic biochemistry platform utilises purified proteins *in vitro* and therefore is not subject to evolutionary pressure to decrease productivity. In principle, any compound can be produced from any starting material without incurring metabolic burden. Synthetic biochemistry can, therefore, be considered ‘evolution-free’ and often results in higher yields of product than can be achieved via *in vivo* production. In our case, we obtained 61 mg L^−1^ raspberry ketone in a batch system, which is approximately 10-fold higher than the highest reported concentration from fermentation. Further improvement could also be achieved with a continuous flow system such as that reported recently by Hold *et al* ^57^ for active provision of substrates and removal of product. Moreover, during our optimisation experiments, we identified that high concentrations of tyrosine as the starting substrate inhibited the overall yield of raspberry ketone. Our study hints that there may be a novel regulation mechanism of the BAS enzyme, possibly allosteric, that prevents over-accumulation of raspberry ketone in its native host. Therefore, synthetic biochemistry can also be a powerful tool for research as well as production of compounds.

Another advantage of the synthetic biochemistry platform is the ability to combine enzymes from different sources with greater ease and less metabolic burden than constructing a strain expressing the whole heterologous pathway. Although we chose to express our enzymes in *E. coli*, in principle pathway variants that are ‘difficult-to-express’ or require post-translational modifications can be expressed and purified in another host, but still mixed with an inexpensive cell-lysate derived from *E. coli* to provide energy metabolism to drive the reaction phase. This hybrid approach is not possible with *in vivo* metabolic engineering where a single production chassis is chosen and its choice may constrain the expression levels of the heterologous pathway or the choice of enzyme source.

Raspberry ketone, although one of the simpler polyketides, still has a complicated biosynthetic pathway that requires ATP, malonyl-coA, and NAD(P)H as cofactors. We showed that each of these could be recycled *in situ* through the application of appropriate cofactor regeneration schemes, but that this must be balanced to avoid the accumulation of side products. In addition, we found that the relative levels of malonyl-CoA, *p*-coumoryl-CoA, and free CoA are a key consideration for maximising yield. The use of an *in vitro* platform allows precise molecular control over the concentration of enzymes, which would be difficult to achieve *in vivo* with the currently available gene expression tools.

To achieve a fine-tuned *in vitro* performance, we have also developed a novel protein-ligand binding fluorescence sensor for direct detection of polyketide synthesis activity and used it to maximise the accumulation of the common intermediates of the raspberry ketone and curcumin biosynthesis pathways. This method could be extended for optimisation of biosynthesis of related polyketides or increase flux towards the malonyl-CoA pool *in vivo*, but will require the inactivation of the CurA reductase activity through point mutations. Given the structural conservation of reductase enzymes, a similar strategy could also be used to develop biosensors for other polyketides in the benzylaklaloid family or to screen variants of the TAL, PCL, or MatB enzymes in order to further increase the yields of raspberry ketone in our synthetic biochemistry platform.

We also demonstrated that raspberry ketone can be synthesised from either tyrosine (the natural substrate *in* vivo), or through a hybrid chemoenzymatic process where HBA is synthesised in one chemical synthesis step, followed by enzymatic conversion to raspberry ketone. *p*-coumarate is also an abundant renewable substrate that can be derived from lignin, so is another plausible starting substrate. The chemoenzymatic approach to high-yield raspberry ketone synthesis provides greater simplicity and lower cost. This method is scalable *in vitro* and potentially also as a microbial fermentation route, potentially providing the most direct method for obtaining industrial yields of raspberry ketone.

Raspberry ketone reductase activity utilises NADPH as a co-substrate which in terms of scalable production is a costly co-factor. To further decrease the cost, we relaxed the cofactor specificity of RKS enzyme to tolerate NADH. A number of studies have highlighted that NADPH-dependent reductases can be engineered to utilise the more stable and inexpensive NADH or biomimetic analogs ^47–50^. As a generalised approach, the structure-guided design of NAD(P)H enzymes can be rationalised by altering the affinity for the ribose 5’-phosphate (NADPH) or hydroxyl (NADH) group ^51,52^. In the case of the RKS enzyme, position G191 provides flexible control for engineering relaxed cofactor specificity. Its coupled use with an inexpensive phosphate donor via the phosphite dehydrogenase ^55^ thus affords a low-cost route to raspberry ketone synthesis *in vitro* with NADH using any of the three starting substrates described.

Approximately 80% of the fine chemical market that is used for cosmetics and food additives, is currently produced by oil derived chemical synthesis and thus approved for use if declared as “nature-identical” ^59^. For fine chemicals that are extracted naturally from food sources that require large agricultural landmasses, therefore, potentially it is far simpler and more sustainable to engineer greener alternative biocatalytic platforms, either from the use of engineered plants and microbes or through *in vitro* isolated enzymes. Here, using raspberry ketone as a model pathway, we have demonstrated how an apparently non-productive enzyme pathway can be engineered to a high level of performance from outside of the cell. Essential to this process is the ability to fine tune enzyme pathways in completion, rather than as individual uncoupled kinetics, since shared resources (CoA, ATP) can impact overall yields. In summary, synthetic biochemistry provides an expandable opportunity to design synthetic enzyme ensembles in unison with cofactor availability. We have applied this rational to optimise raspberry ketone, a small molecule that is not easily obtained from within engineered living cells.

## Materials and Methods

### Molecular biology and protein expression

Routine molecular biology was performed as described previously by our previous research ^60^. For the synthetic raspberry ketone pathway the following enzymes were selected - *Rhodotorula glutinis* TAL ^37^, *Arabidopsis thaliana* PCL ^35,38^, *Rhodopseudomonas palustris* MatB ^61^, *Rheum palmatum* BAS ^25,28^ and *Rheum idaeus* RKS from ^36^. The genes encoding these enzymes were synthesised by ThermoFisher Scientific and codon optimised for *E. coli* K12 expression with compatibility for EcoFlex cloning ^60^. PtxD was used for NAD(P)H recycling from phosphite ^55^, whilst ATP regeneration was provided from PEP and rabbit pyruvate kinase (Sigma, UK). All oligonucleotides, plasmids and synthetic DNA sequences are listed in the supporting information. Sequencing was performed by Eurofins, Germany.

### Golden Gate Mutagenesis

We developed a new protocol for mutagenesis based on Golden Gate cloning. Forward and reverse primers were designed for inverse PCR of the plasmid (pTU1-A-T7His-RKS-Bba_B0015) to incorporate a BsaI site, which is routinely used in Golden Gate cloning for directional assembly. After digestion with BsaI, the restriction sites are removed, leaving a complementary 4 bp overhang, which is designed to incorporate the mutation site. This anneals and ligates to provide the desired mutation. PCR was performed with the Q5 polymerase (NEB, UK) with 3% DMSO using the manufacturers standard guidelines. PCR was run on a 1% agarose gel and the bands were excised and gel purified with a QIAquick Gel Extraction Kit. 20 ng of PCR product was then digest and ligated in a one-pot reaction with 1 × DNA ligase buffer (Promega), 1 unit of Bsal-HF, 5 units of T4 ligase and 1 unit of DpnI. The reaction was run for 15 cycles at 37°C for 5 min and 16°C for 10 min, followed by a further incubation at 50°C for 5 min and 80°C for 5 min. 20 μL of DH10β competent cells were transformed with 2μL of ligation mix and plated onto 100 μg mL^−1^ carbenicillin plates. Single colonies were then sequence verified for incorporation of the mutation site.

### Protein expression and purification

His_6_-tagged recombinant TAL, PCL, BAS, MatB and RKS were over-produced in *E. coli* BL21-Gold (DE3) grown at 37°C, 200 rpm in 2YT medium with 100 μg/ml ampicillin until an OD_600_ of 0.6 was reached. Cells were induced with 0.4 mM IPTG and grown overnight at 21°C at 200 rpm. Cell were collected by centrifugation at 6,000 × *g*, 4°C for 20 min, then re-suspended in binding buffer (20 mM Tris-HCl pH8, 500 mM NaCl, 5 mM imidazole) and lysed by sonication. Cell-lysates were clarified with centrifugation at 45,000 × *g*, 4°C for 20 min and purified by gravity flow using Ni-NTA agarose (Generon). His_6_-tagged proteins were washed with increasing concentrations of imidazole (5, 30 and 70 mM) in 20 mM Tris-HCl pH 8, 500 mM NaCl, before elution at 400 mM imidazole. Purified protein were then dialysed (MWCO 10,000) into 2 litres of 20 mM HEPES pH 7.5, 100 mM NaCl (Buffer A) at 4°C for 6 hours. The enzymes were found to be soluble and active in a range of standard buffers including HEPES (pH 7.5) and Tris-HCL pH 8.0-9.5. Additionally, all enzymes were stable for long-term storage at −80°C with 15% (v/v) glycerol.

### Chemical synthesis of HBA

HBA, also referred to as 4-(4-hydroxyphenyl)-buten-2-one was synthesised by a crossed aldol condensation as previously described ^62^. Further details and NMR spectra (Supplementary Figure S10) are provided in the supplementary data.

### LC-MS of raspberry ketone and pathway intermediates

50 μL samples of time-course reaction in triplicate were removed and inactivated with 450 μL of 1% HCl. Samples were centrifuged at 13,000 rpm for 10 min at room temperature. The supernatant was directly analysed by liquid-chromatography mass spectrometry (LC-MS), performed with an Agilent 1290 Infinity system with an online diode array detector in combination with a Bruker 6500 quadruple time-of-flight (Q-ToF) mass spectrometer. An Agilent Extend-C18 2.1 × 50mm (1.8 μm particle size) column was used at a temperature of 25 °C with a buffer flow rate of 0.2 ml^−1^ min^−1^. LC was performed with a linear gradient of buffer A (0.1% formic acid) and buffer B (0.1% formic acid in acetonitrile). Separation was achieved using 5% buffer B for 2 min, followed by a linear gradient to 50% buffer B from 2-9 min, which was held at 50% buffer B from 9-10 min. Spectra were recorded between a mass range of 90-1000 *m/z* at a rate of 3 spectra per second. Standards were prepared and calibration curves for the intermediates tyrosine, *p*-coumaric acid, HBA and raspberry ketone were derived. Quantitation was based on the MS peak area of precursor or fragment ion in comparison to the analytical standards. Under the conditions used, raspberry ketone is detected as a sodium adduct [M+Na^+^]^+^ or as a diagnostic fragment ion at *m/z*= 107.49, corresponding to C_7_H_7_O. For the standards in solvent, good linearity (R^2^>0.99) was achieved over the range of 0.3 to 30 pmol on column. The lower limit of quantitation was set at 0.3 pmol. Samples that were below this limit were repeated by increasing the injection volume to 1 μl. Due to a lack of a analytical standard and poor separation, *p*-coumoryl-CoA was not quantified.

### Enzyme kinetics

RKS and mutants were purified to homogeneity using nickel IMAC and buffer exchanged into Buffer A. Steady-state kinetics were monitored on a Clariostar (BMG Lifetech) plate-reader monitoring absorbance at 340 nm following the reduction of NAD(P)H to NAD(P)*+* with either 4-hydroxyphenyl-3-butan-2-one (hydroxybenzaldehyde) or phenyl-3-butan-2-one (benzaldehyde) as substrates. Assays were performed in triplicate at 30°C in 0.1 M potassium phosphate pH 6.4.

### RKS crystallisation and structure determination

IMAC purified RKS was dialysed for 4 hours in 20 mM Tris-HCL pH 8.0 and 200 mM NaCl. Pure fraction concentrated with a 10,000 MWCO centrifugation concentrator (Amicon) and then run on analytical gel filtration in the same buffer. Purity was assessed by SDS-PAGE and the concentration was determined by A_280_ measurement using an extinction coefficient of 44,030 M^−1^ cm^−1^. RKS was concentrated to 10 mg mL^−1^ and screened in a range of crystallisation conditions using 300 nL drops containing either a ratio of 1:2 or 2:1 of protein and reservoir buffer. Crystals of N-terminally His_6_-tagged RKS were obtained by sitting drop vapour diffusion at 20°C after ~3 days of incubation in 0.1 M MES/imidazole pH 6.3, 11% (w/v) PEG 550 MME and 5% (w/v) PEG 20K, 20 mM of amino acid mixture (Molecular dimensions Morpheus system) with 1 mM NADPH. Single cube-shaped crystals grew within 1 week. Native crystals with NADPH bound were soaked in cryoprotectant containing 1 mM HBA, 1 mM NADPH and 20% glycerol. A native dataset of 1800 frames was collected remotely at the I04 beamline (Diamond Light Source, Didcot, Oxfordshire) from a single crystal diffracting up to ~1.5 Å. The crystal belonged to space group P1 (Supplementary Table S1). Further details on structure determination are provided in the supplementary text. The atomic coordinates and structure factors (codes: 6EOW for the ternary structure) has been deposited in the Protein Data Bank.

## Acknowledgements

We would like to thank Dr Tobias Erb (Max Planck Institute) for constructive comments on the preliminary data from this project. We would also like to thank Dr Marc Morgan, Dr Inmaculada Pérez-Dorato and Professor Angelika Gründling for project support concerning the crystallography of RKS, and all the staff in the beam line I04 at the Diamond Light Source (Didcot, Oxfordshire, UK) where the crystallographic datasets were collected. We would also like to thank the EPSRC for funding to SM (EP/K038648/1 - Frontier Engineering).

